# Chronic striatal cholinergic interneuron excitation induces clinically-relevant dystonic behavior in mice

**DOI:** 10.1101/2023.07.19.549778

**Authors:** Kat Gemperli, Xinguo Lu, Keerthana Chintalapati, Alyssa Rust, Rishabh Bajpai, Nathan Suh, Joanna Blackburn, Rose Gelineau-Morel, Michael C. Kruer, Dararat Mingbundersuk, Jennifer O’Malley, Laura Tochen, Jeff Waugh, Steve Wu, Timothy Feyma, Joel Perlmutter, Steven Mennerick, Jordan McCall, Bhooma R. Aravamuthan

## Abstract

Dystonia is common, debilitating, often medically refractory, and difficult to diagnose. The gold standard for both clinical and mouse model dystonia evaluation is subjective assessment, ideally by expert consensus. However, this subjectivity makes translational quantification of clinically-relevant dystonia metrics across species nearly impossible. Many mouse models of genetic dystonias display abnormal striatal cholinergic interneuron excitation, but few display subjectively dystonic features. Therefore, whether striatal cholinergic interneuron pathology causes dystonia remains unknown. To address these critical limitations, we first demonstrate that objectively quantifiable leg adduction variability correlates with leg dystonia severity in people. We then show that chemogenetic excitation of striatal cholinergic interneurons in mice causes comparable leg adduction variability in mice. This clinically-relevant dystonic behavior in mice does not occur with acute excitation, but rather develops after 14 days of ongoing striatal cholinergic interneuron excitation. This requirement for prolonged excitation recapitulates the clinically observed phenomena of a delay between an inciting brain injury and subsequent dystonia manifestation and demonstrates a causative link between chronic striatal cholinergic interneuron excitation and clinically-relevant dystonic behavior in mice. Therefore, these results support targeting striatal ChIs for dystonia drug development and suggests early treatment in the window following injury but prior to dystonia onset.

**One Sentence Summary:** Chronic excitation of dorsal striatal cholinergic interneuron causes clinically-relevant dystonic phenotypes in mice

## INTRODUCTION

Dystonia is a functionally debilitating, painful, and often medically refractory movement disorder characterized by overflow muscle activation triggered by voluntary movement and is, by definition, a variable phenomenon.(*1–5*) Mouse models of rare, monogenic, and progressive etiologies of dystonia have provided some insights into underlying pathophysiology, particularly of early-onset dystonias (onset during childhood or early adulthood). In particular, many of these studies have demonstrated abnormal excitability of striatal cholinergic interneurons (ChIs), suggesting a potential pathophysiologic link.(*6–9*) However, it is unclear if excitation of striatal ChIs can independently cause a dystonic phenotype. Optogenetic excitation of striatal ChIs for several minutes does not produce classic dystonic behaviors in mice (*10*), which are primarily characterized as clasping behaviors observed during tail suspension.(*11–13*)

Although early-onset dystonia pathophysiology has been most commonly studied in the context of rare monogenic progressive etiologies,(*6*, *7*, *9*, *10*, *14–23*) the most common cause of early-onset dystonia is neonatal brain injury producing cerebral palsy (CP).(*24–27*) In CP, dystonia appears after a temporal delay of months to years after the injury.(*3*, *28*) It is unclear whether this delay following injury is necessary for dystonia to develop. However, conceptualization of dystonia as a circuit disorder has led to the hypothesis that underlying circuit pathology must persist for a threshold duration of time to yield dystonia.(*29*) For example, long-term striatal ChI excitation beyond what has been assessed in optogenetic studies potentially could yield dystonic behavior in mice.

The definition of dystonic behavior in mice, however, is challenging. Although classically characterized as clasping, several caveats preclude direct translational relevance of this mouse behavior to people with dystonia. First, clasping is a binary phenomenon. Although, the length of time that clasping lasts can be quantified during a given behavioral task, no agreed upon criteria exist to differentiate less versus more severe clasping. Second, clasping or other previously defined behaviors have not been identified in many genetic mouse models of dystonia that recapitulate clinically relevant mutations (e.g. the mouse model of the *Tor1a* GAG deletion that is associated with DYT1 dystonia).(*11–13*) Finally, and perhaps most importantly, clasping is not a motor behavior directly related to dystonia in people.(*30–32*) Electromyographic measures of antagonist muscle co-contraction have also been used to quantify dystonia in rodents.(*19*, *33*, *34*). However, antagonist muscle co-contraction is not specific to dystonia alone and can be seen associated with other motor symptoms like spasticity,(*35*, *36*) which often co-occurs with dystonia and requires a different treatment paradigm.(*37*, *38*) To quantify the most clinically relevant dystonic behaviors in mice, it is critical to first codify behaviors specifically characteristic of dystonia in people.

Clinically, the gold standard for identification of dystonia relies on expert consensus-based assessment of physical exam videos.(*31*, *32*) The specific features governing a clinical dystonic movement have not been comprehensively identified. Recent work has demonstrated that dystonia in the legs, the most common site of dystonia in people with CP, primarily manifests during gait as quantifiably variable leg adduction.(*32*, *37*) It is unclear whether leg adduction variability tracks with dystonia severity during other tasks as well. If so, it may be possible that leg adduction variability in mice during the tail suspension task may be a more clinically-relevant dystonia marker than current clasping metrics.

Here, we demonstrate that chronic striatal ChI excitation can yield such clinically-relevant dystonic behavior in mice. To do so, we first show that variable leg adduction correlates with dystonia severity in people with CP as they are doing seated tasks which do not otherwise engage the legs (similar to how the mouse tail suspension task should not perpetually engage the hindlimbs). We next show that chronic chemogenetic striatal ChI excitation in mice over two weeks causes hindlimb adduction variability during tail suspension. These results demonstrate, for the first time, that striatal ChI excitation can cause clinically-relevant dystonic behavior in mice and that these dystonic behaviors only emerge after a prolonged period of excitation (i.e. a delay). These results support treatment development focused on decreasing striatal cholinergic tone in people with dystonia and support current clinical hypotheses suggesting it is most advantageous to treat dystonia as early as possible.

## RESULTS

### Leg dystonia severity is primarily associated with leg adduction variability in people with CP

We previously demonstrated in people with CP that leg adduction variability correlates linearly with expert assessed leg dystonia severity during gait.(*32*) To determine the robustness of this association across tasks, including tasks that do not directly involve the legs, we had eight pediatric movement disorders neurologists from eight different institutions rate leg dystonia in videos of people with CP while they were engaging in a seated hand open/close task. Dystonia severity was rated using the Global Dystonia Rating Scale (GDRS), which has high inter-rater reliability for leg dystonia severity assessment.(*32*, *39*)

Experts noted that 104/193 people assessed (demographic characteristics in Supplemental Table 1) had some degree of leg dystonia, comparable to published rates of dystonia in people with CP.(*31*, *32*, *37*) 52/104 had mild dystonia (GDRS 1 to <4), 25/104 had moderate dystonia (GDRS 4 to <7), and 27/104 had severe dystonia (GDRS 7 or more). Using DeepLabCut, we determined the 2D coordinates of the knees, ankles, and toes of each subject in each video frame and then calculated the leg adduction amplitude and variability metrics that we have previously shown correlate with leg dystonia severity during gait. Almost all assessed metrics of leg adduction amplitude and variability were significantly greater in those with leg dystonia compared to those without (t-test, p<0.05) (Fig. 1). Almost all of these metrics also correlated with GDRS leg dystonia severity score (Pearson correlation, p<0.01) (Supplemental Figure 1).

**Fig. 1.**
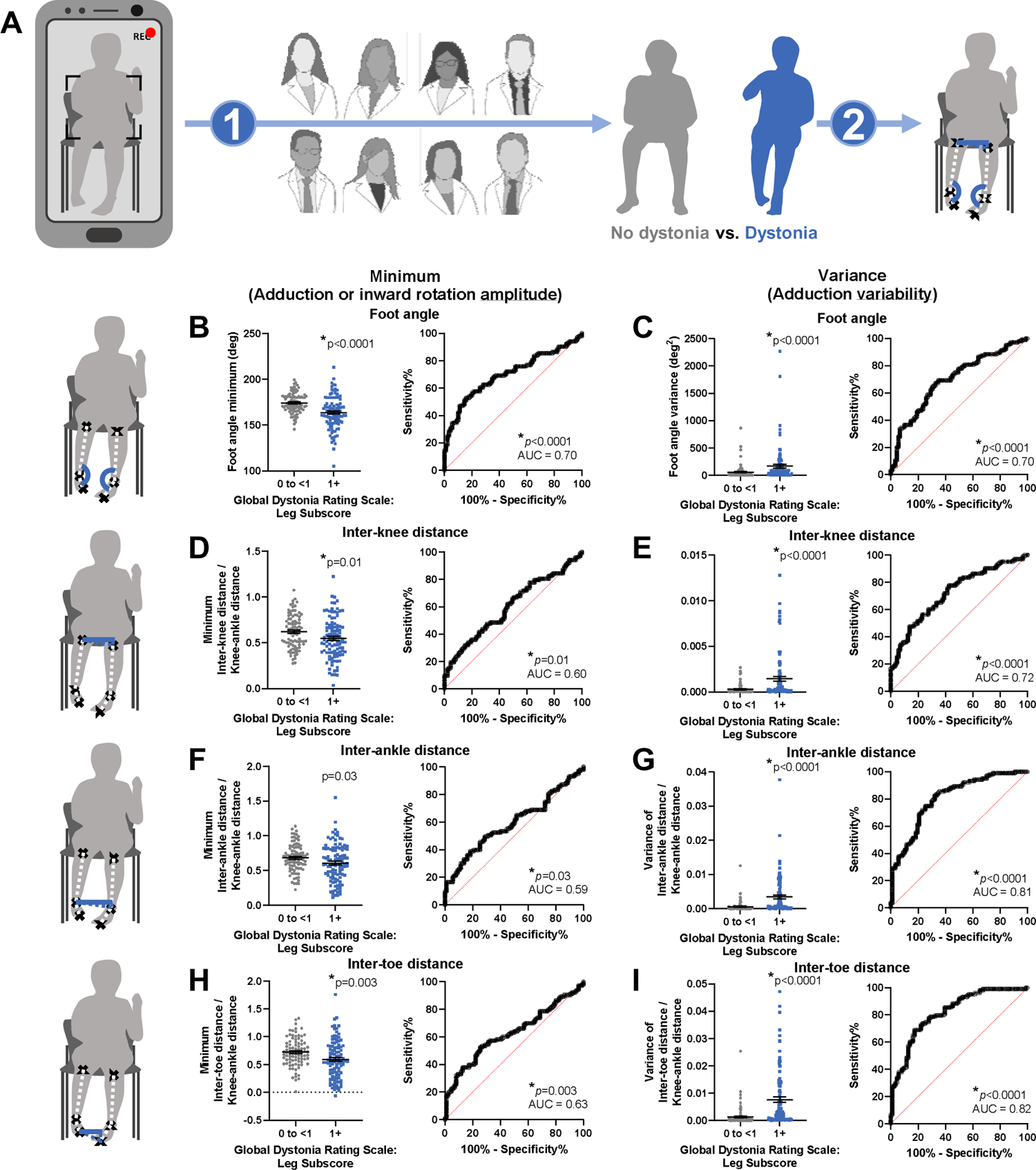
Differences in leg adduction variability and amplitude between people with and without leg dystonia. **A)** Overall experimental paradigm: Videos of seated subjects doing a hand open-close task are recorded, assessed by eight pediatric movement disorders physicians for dystonia using the Global Dystonia Rating Scale (GDRS), and then assessed for quantifiable metrics of experts’ dystonia assessments. **B-I)** Metrics of leg adduction amplitude or minima (left two columns) and variability or variances (right two columns) are compared between people with CP and leg dystonia (blue, n=103, average GDRS of 1 or more) and people with CP but without leg dystonia (grey, n=90, average GDRS of 0 to <1). Metrics are the minima and variances of the foot angle in the coronal plane (angle between the knee, ankle, and toe labels) (**B&C**), inter-knee distance (**D&E**), inter-ankle distance (**F&G**), and inter-toe distance (**H&I**) with distance metrics assessed exclusively along an X-axis parallel to the floor. Comparisons between people with CP with and without dystonia are by t-test with receiver operator characteristic curves (AUC – area under the curve) showing the sensitivity and specificity of each metric for expert-assessed dystonia, *p<0.05.

To determine which quantified metrics served as the best objective measures of dystonia, we used multiple linear regression models, receiver-operator-characteristic curves, and multiple feature selection methods. These all showed that leg adduction variability related to dystonia severity much better than did leg adduction amplitude. A multiple linear regression model relating dystonia severity to only the four variability metrics had an adjusted R^2^ of 0.546 (F 57.6, p<0.001), whereas a model including just the four amplitude metrics had a lower adjusted R^2^ of 0.268 (F 18.2, p<0.001). Including all eight assessed metrics of variability and amplitude in a multiple linear regression only marginally improved the adjusted R^2^ to 0.584 (F 34.0, p<0.001) compared to only including the variability metrics. Furthermore, metrics related to variability were best able to discriminate between dystonia and no dystonia as indicated by receiver-operator-characteristic curves, with three of the four variability metrics demonstrating the highest areas under the curves (Fig. 1). Finally, we used 7 filter, wrapper, and embedded feature selection methods to either assess how well variability or amplitude metrics can discriminate between the presence or absence of dystonia or predict dystonia severity. All 7 of these methods selected variability metrics more often than amplitude metrics as the primary contributors to models either discriminating between the presence or absence of dystonia or models predicting dystonia severity (Supplemental Table 2). Taken together, these data suggest that leg amplitude variability in people with CP relates to dystonia severity better than amplitude metrics.

### Leg adduction variability increases with chronic striatal ChI excitation in mice

We next sought to determine how these clinically-derived metrics of expert-assessed leg dystonia severity could be leveraged to understand dystonia pathophysiology in mice (Fig. 2A). Striatal ChIs are abnormally excitable in genetic mouse models of dystonia,(*8*, *40*) are increased in a rat model of cerebral palsy and dystonia,(*41*) and are preferentially lost in the dorsal striatum in a genetic mouse model of DYT1.(*42*, *43*) To determine whether chronic striatal ChI excitation could cause clinically-relevant dystonic behavior in mice, we targeted expression of excitatory Gq designer receptors exclusively activated by designer drugs (DREADDs) to the dorsal striatum using stereotaxic injections of AAV8-hSyn-DIO-hM3d(Gq)-mCherry. The experimental group (ChAT-Cre positive mice on a C57/Bl6J background) expressed these excitatory DREADDs in ChIs in the dorsal striatum while the control group (ChAT-Cre negative littermates) did not. We saw that DREADD expression co-localized to ChAT expression in the dorsal striatum in the experimental group (Fig. 2B), but that a relatively small percentage of the total striatal ChIs in any given slice were labeled (2-18% of striatal ChI cell bodies visible on any slice between 145 and 945 μm anterior to bregma) (Fig. 2C).

**Fig. 2.**
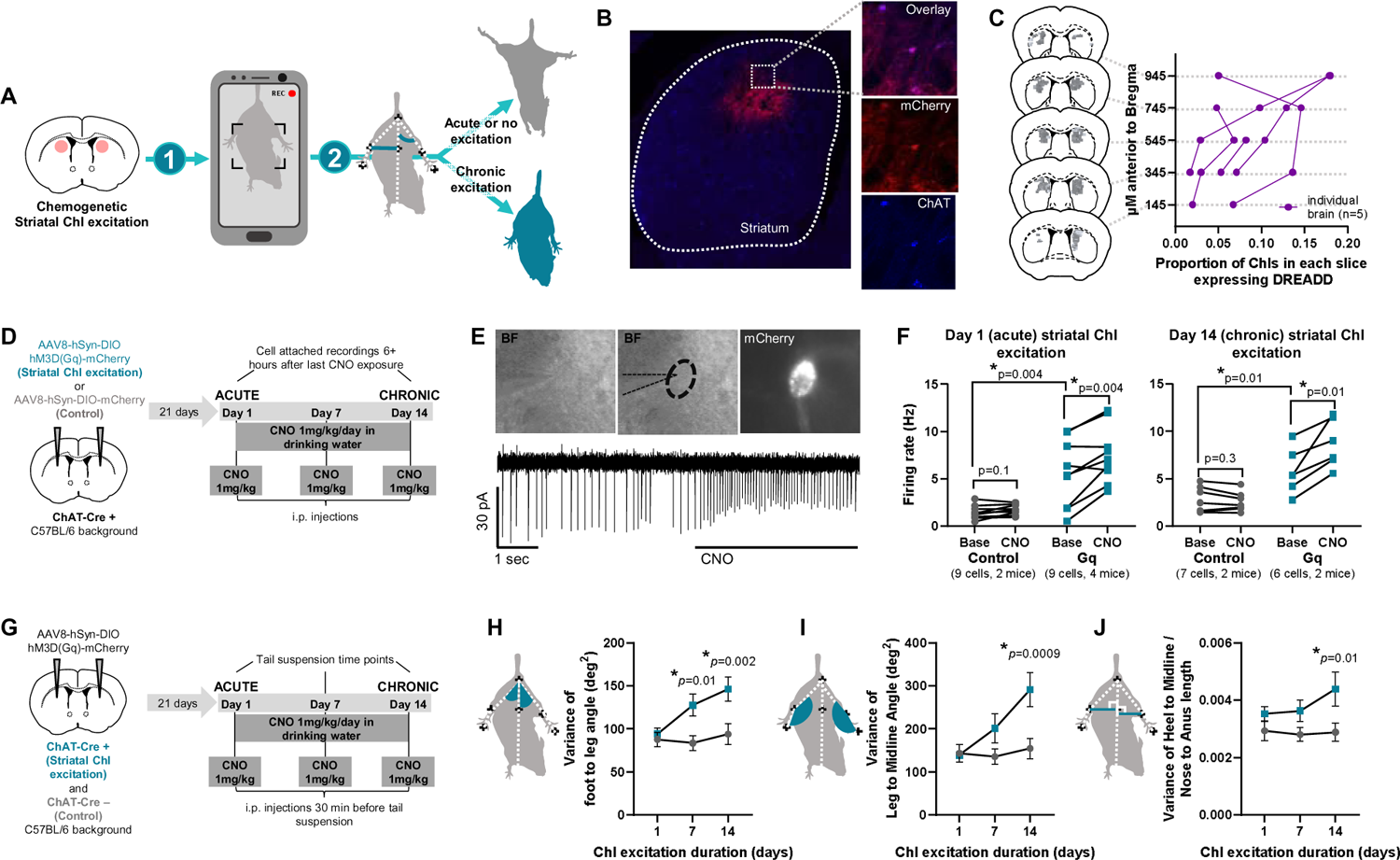
Leg adduction variability in mice in response to 14 days of dorsal striatal ChI excitation. **A)** Overall experimental paradigm: Striatal cholinergic interneurons (ChIs) are chemogenetically excited in mice over 14 days, during which videos during a tail suspension task are recorded. These tail suspension videos are assessed for quantitative metrics of dystonia validated in people with CP. Differences in these clinically-derived dystonia metrics are compared between mice who do and do not undergo chronic striatal ChI excitation. **B)** Immunohistochemical staining of striatal slices for choline acetyl-transferase (ChAT), mCherry (co-expressed with the Gq DREAD), and overlay images show that ChAT and mCherry co-express in the same neurons. **C)** Mapping of the location and proportion of striatal ChIs expressing the Gq DREADD between 145-945 μm anterior to Bregma. **D)** Experimental paradigm showing generation of mice expressing the mCherry fluorescent tag in striatal ChIs (grey, control) and animals additionally expressing the Gq DREADD (teal, experimental) and the shared 14 day CNO exposure protocol and cell-attached recording timepoints. **E)** Top: Example of cell-attached patching of a striatal ChI under brightfield (BF) and fluorescence to visualize the mCherry fluorescent tag. Bottom: An example tracing of striatal ChI firing and response to bath-applied CNO (in a neuron co-expressing the Gq DREADD). **F)** Striatal ChI firing rates and responses to bath-applied CNO determined with cell-attached recordings in mice on Day 1 of the CNO exposure paradigm and Day 14 of the CNO exposure paradigm, with baseline firing rates recorded at least 6 hours after the last CNO exposure. Paired t-tests for comparing pre- and post-CNO firing rates; unpaired t-tests for comparing baseline firing rates between groups, *p<0.05. **G)** Experimental paradigm showing generation of mice expressing the Gq DREADD (teal, ChAT-Cre+, experimental) or not (grey, ChAT-Cre-, control) and the shared 14 day CNO exposure and tail suspension recording timepoints. **H-J)** Leg adduction variability metrics during tail suspension over the 14-day striatal ChI excitation period are shown. Variances of the foot angle between the hindpaws and legs (**H**), leg angle between the leg and the mouse midline (**I**), and distance between the heels and midline (**J**) are shown. Comparisons between mice that did and did not undergo ChI excitation are by repeated measures two-way ANOVA with Bonferroni correction for multiple comparisons, *p<0.05.

To validate that striatal ChIs were, in fact, being excited with the DREADD agonist clozapine-N-oxide (CNO), we made cell attached recordings from striatal ChIs expressing either the excitatory Gq DREADD or a fluorescent mCherry reporter (Fig. 2D-E). As demonstrated previously, striatal ChIs expressing the Gq DREADD had a significantly higher baseline firing rate (t-test, p<0.05) which was potentiated with CNO (paired t-test, p<0.05) (Fig. 2F).(*44*) This was observed in mice on day 1 and day 14 of ongoing CNO exposure (Fig. 2F), demonstrating that mice in the experimental group were undergoing chronic striatal ChI excitation during the behavioral paradigm, while mice in the control group were not.

Despite targeting only a small proportion of striatal ChIs, sustained excitation of these neurons over 14 days with repeated administration of CNO increased leg adduction variability during tail suspension (Fig. 2G-H). Foot angle variance, leg angle variance, and variance of the distance between the hindpaw heel and trunk midline increased over time and significantly differed between the control (n=17) and experimental groups (n=17) by day 14 of ongoing striatal ChI excitation (Fig. 2H, two-way repeated measures ANOVA, p<0.05). However, there was no significant difference between groups on day 1 of behavioral assessment, suggesting that chronic striatal ChI excitation over a period of several days is required to yield this clinically-relevant dystonic behavior. There was no significant effect on leg adduction amplitude during tail suspension (Supplemental Figure 2A). Importantly, the increase in leg adduction variability was not solely due to increased leg movement. Total movement of the hindpaw (measured at the heel and toe relative to the trunk midpoint) did not differ significantly between groups across the whole 14 day span (Supplemental Figure 2B).

## DISCUSSION

Dystonia is a common, debilitating, and often medically refractory movement disorder.(*3–5*) The development of new treatments for dystonia has been limited, in part, by the difficulty in establishing clinically-relevant but quantifiable metrics of dystonia that can be assessed across species. We have demonstrated that expert assessment of leg dystonia severity primarily relates to quantifiable leg adduction variability and that this leg adduction variability emerges after chronic striatal ChI excitation in mice. Therefore, we demonstrated a potentially causative link between chronic striatal ChI excitation and clinical relevant dystonic behavior.

Abnormal striatal ChI excitability has been a unifying feature of many mouse models of genetic dystonias.(*8*, *40*) The loss of these neurons in the dorsal striatum temporally relates to the onset of dystonic symptoms in a mouse model of DYT1.(*43*) Yet, a causative link remained elusive, noting that previous work has examined behavioral outcomes after exciting striatal ChIs in mice for only minutes.(*45*) Furthermore, the behavioral outcomes that have been classically assessed as dystonic in mice have no direct behavioral correlate in people.(*30–32*)

We demonstrate that striatal ChI excitation can cause clinically-relevant dystonic behavior in mice, and show that this excitation has to occur over a prolonged period of time (at least 14 days of continuous excitation). This finding has immediate clinical implications. It has long been established that dystonia emerges following a delay of months to years after a brain injury(*3*, *28*) and that, in the context of CP which is the most common cause of dystonia in childhood, that 50-80% of people have some degree of dystonia.(*31*, *32*, *37*, *46*) These results potentially suggest that reducing striatal ChI excitation in the period following brain injury but before the emergence of dystonia, may mitigate or possibly prevent dystonia manifestations. Furthermore, people with dystonia respond best to surgical treatments (i.e., deep brain stimulation) if their dystonia is treated early.(*47*, *48*) It may also be possible that preventative measures could be tested for presymptomatic people at high risk for dystonia (e.g., in children in the months to years following neonatal brain injury). At the very least, these results advocate for high vigilance and early diagnosis of dystonia so that early treatment (pharmacologic or otherwise) can be considered.

Our approach also may facilitate the diagnosis of leg dystonia in people with CP. Dystonia most commonly presents in the legs in people with CP(*37*) and typically begins in the legs in people with the most common monogenic form of dystonia (DYT1).(*5*, *49–51*) The legs are also where dystonia is most commonly underdiagnosed.(*38*) We have previously established that leg adduction variability relates to leg dystonia severity during gait in people with CP.(*31*, *32*) However, using gait alone as the task to facilitate dystonia identification requires assessment outside of a standard clinical exam room and also precludes the assessment of non-ambulatory individuals. Here, we show that leg adduction variability can still be used to assess individuals for leg dystonia during seated upper extremity tasks. The immediate clinical implications of these results are two-fold. First, leg adduction variability can be considered as a robust metric for leg dystonia assessment as it has been demonstrated across two large cohorts of individuals across two different motor tasks.(*31*, *32*) Second, either qualitative or quantitative assessment of leg adduction variability in the clinical setting using gait or a seated hand open/close task could help facilitate clinical dystonia diagnosis.

This study only established clinically-relevant metrics of dystonia in people with CP. Although CP is the most common cause of early-onset dystonia with heterogenous etiologies,(*24*, *26*, *27*), CP may not necessarily represent the way dystonia manifests across myriad rarer conditions. A similar approach should be tested in people with rarer etiologies of dystonia. Additionally, though we assessed mice during the task typically used to assess for dystonia (tail suspension), assessing mice for clinically-derived dystonia metrics across other tasks would also be worthwhile and would likely further establish the translational relevance of these metrics. Finally, we note that striatal ChI excitation in mice expressing Gq DREADDs appeared to occur at baseline even prior to CNO administration, as demonstrated previously.(*44*) This suggests that the time window of ongoing striatal ChI excitation required to yield dystonia is *at least* 14 days, but may be longer (given that we waited 21 days to allow DREADD expression before beginning to assess behavior). Future work could assess whether exogenous expression of Gq DREADDs in striatal ChIs in isolation, without administration of a DREADD agonist, would result in dystonic behavior in mice. However, even noting that striatal ChIs increase their firing rate with expression of the Gq DREADD alone, these data support the conclusion that chronic striatal ChI excitation is necessary to cause dystonic behavior in mice, in line with the clinically observed delay between an inciting brain injury and resultant dystonic behavior in people.

Future work should also consider incorporating quantification of leg adduction variability across behavioral tasks when assessing dystonia in mouse models. Though clasping of the fore- or hind-paws during tail suspension has been considered as the canonical metric for dystonia identification, our results show that the clinically-relevant metric is leg adduction variability. Furthermore, we suggest that reverse translational behavioral assessments of qualitative clinical phenomena should begin by first establishing unbiased quantifiable (i.e., not-manually scored) metrics of the behavior in people and then developing ways to assess similar metrics in animal models of disease. This approach may be most likely to yield the greatest translational relevance of animal model data.

## MATERIALS AND METHODS

### Study Design

We hypothesized that chronic striatal ChI excitation would result in clinically-relevant dystonic behavior in mice.

Previous work has shown that leg adduction variability and amplitude during gait is significantly greater in people with leg dystonia compared to people without leg dystonia.(*31*, *32*) Based on the effect sizes from previous work, a sample size of 39 subjects per group would be necessary to detect comparable differences between people with and without dystonia (effect size d = 0.57, 80% power, α 0.05).(*52*) Videos were collected comparably across all subjects. Experts evaluated videos of subjects for dystonia blinded to subject identity (faces blurred) without any other knowledge of the subject except that they had been diagnosed with CP.

Mouse experiments were conducted across two different experimental cohorts (each of which included control and experimental groups). Randomization to control and experimental groups occurred based on genotype within each litter (see “Chemogenetic excitation of striatal ChIs in mice”). Behavioral data collection and initial processing were done by individuals blinded to the experimental status of individual mice (see “Clinical expert-based leg dystonia assessment in human subjects”).

### Human subjects and video collection

All human subject data collection was approved by the Washington University School of Medicine Institutional Review Board.

Subjects with CP were recruited from the Gillette Children’s Cerebral Palsy Center between 1/1/2020 and 12/30/2021. Inclusion criteria were that subjects had to be at least 5 years old (to optimize the likelihood that a child would have the cognitive ability and attention span to reliably complete the task) and that they had to have a clinician-confirmed diagnosis of CP as per the 2006 consensus diagnostic criteria.(*53*) Exclusion criteria were inadequate quality or content of the video recordings (e.g. full body not visible for the task or the person was not able to complete the task during the video recording). Subjects were asked to do an alternating hand open/close task: they sat in a chair facing forward, rested the non-dominant hand on the ipsilateral thigh, and then raised and alternatingly opened and fisted the dominant hand as fast they were able for approximately 5 seconds. They then repeated this task with the non-dominant hand. Videos were recorded with a Panasonic AG-AC160AP camera at 1920 x 1080 pixel resolution.

### Clinical expert-based leg dystonia assessment in human subjects

For video assessment, faces of all subjects were blurred to ensure subject anonymity using ShotCut (Meltytech, LLC). Initially, blurred videos were assessed for leg dystonia using the Global Dystonia Rating Scale (GDRS) by one pediatric movement disorders physician (BRA). The GDRS is a 10-point Likert scale score (0 – no dystonia, 10 – severe dystonia) which has demonstrated good inter-rater reliability for assessment of leg dystonia in people with CP.(*32*, *39*) These preliminary GDRS scores were not included in the final analysis or shared with other video reviewers but were used to roughly stratify videos across a possible range of dystonia severities. This rough stratification was used to create three consensus-building data sets of 8 videos each across a putatively broad range of dystonia severities (including videos where dystonia was absent based on these initial ratings). The three video sets were presented to 8 pediatric movement disorders physicians with expertise in CP practicing at eight different institutions across the United States (JB, RGM, MK, DM, JOM, LT, JW, SW). This group of eight reviewers met for three 1-hour Zoom-based consensus-building discussions to ensure shared understanding and application of the GDRS for leg dystonia rating in CP (Zoom Video Communications, Inc.). During each 1 hour discussion session, each of the 8 videos were discussed as follows: 1) The video was played on loop while it was being discussed; 2) When ready, pre-assigned primary and secondary discussants described how they would approach leg dystonia assessment in the video; 3) The whole group was invited to discuss their assessments of the video, as desired; 4) Each of the 8 experts independently entered their leg dystonia GDRS scores for the video into a REDCap survey (REDCap 12.4.31).

After 24 videos were assessed across these three consensus-building sessions, the remaining videos were independently assessed for leg dystonia and scored using the GDRS by two of the eight experts, again with scores entered via a REDCap survey. Each expert was assigned videos across a range of possible dystonia severities. For any videos where the total leg dystonia GDRS score differed by more than 6 between the two reviewers, the whole group of eight experts was asked to assess the video including re-assessment by the two original reviewers. The average score across the two reviewers, or all across all eight reviewers for the consensus building video data set or for videos with scoring discrepancies, were used for final analysis.

### Mouse tail suspension task and video collection

All experiments and procedures were approved by the Institutional Animal Care and Use Committee of Washington University School of Medicine in accordance with National Institutes of Health guidelines.

Mice were from litters resultant from breeding Choline acetyl-transferase-Cre (ChAT-Cre, Jackson Laboratory Strain #006410) mice with wild-type C57/Bl6J (Jackson Laboratory Strain #000664) mice. Both sexes were used for all experiments. Mice were held 1 inch above the tail base and suspended 1 foot above a table surface in front of a solid colored background for 1 minute. Videos of the ventral side of the mouse were recorded using a Google Pixel 2 (Google, Mountain View, CA, USA) on a tripod placed 6 inches in front of the mouse. Videos were recorded at 120 frames per second at 1980 x 1020 pixel resolution.

### Video-based pose estimation and calculation of clinically-relevant dystonia metrics

DeepLabCut was used to estimate the positioning of the legs in human hand open/close videos and of the hindlimbs in mouse tail suspension videos. Noting that variable leg adduction has been described as a key feature of leg dystonia during gait in people with CP, leg and hindlimb features that allow for quantification of leg adduction variability were labeled.

In human subjects, the following points were determined across all videos for the right and left leg: midpoint of the patella, midpoint between the medial and lateral malleolus, and tip of the third toe. These points were manually labeled on 10 frames extracted from each video and used to train a ResNet-50-based neural network for 1,030,000 iterations with 95% of frames used for training and 5% of frames used for testing the network to yield less than a 5 pixel train error and 5 pixel test error. 10 videos labeled using the trained network were visually examined in their entirety to ensure the network was appropriately labeling the desired points.

In mice, the following points were determined across all videos for the right and left hindlimbs: hindpaw heel and hindpaw mid-toe. Given that a point analogous to the patella is not readily visualizable in videos of mice, the following mouse midline features were labeled to allow for assessment of hindlimb adduction variability and amplitude: anus, the trunk midpoint between the two hindlimbs, and the nose tip. These points were manually labeled on 10 frames extracted from 30 videos taken on different experimental days and from different experimental cohorts. These manually labeled frames were used to train another network as described above. Points labeled using these trained networks were conditioned at a p-cutoff of 0.99 to ensure only accurately labeled points were used for further analysis.

These leg adduction variability and amplitude metrics were calculated across all videos using custom written code in MATLAB (R2021 Mathworks Inc.) for both human subjects and mice. The MATLAB code and example trimmed labeled videos are available in the Supplemental Materials and Methods and as Movies S1-S5. All supplemental files including full-length videos and trained DeepLabCut labeling models are available here: https://wustl.box.com/s/07agml8jmw681et1bnsk2j4atghy5j9l.

### Chemogenetic excitation of striatal ChIs in mice

Mice were genotyped via tail snip at post-natal day 28. All mice underwent the following surgical and chemogenetic excitation protocol with experimental groups decided by mouse genotype (ChAT-Cre positive mice in the experimental group were compared to ChAT-Cre negative controls). Between 5-6 months of age, mice were stereotaxically injected under isoflurane anesthesia (1% v/v in oxygen, 1 L/min) with 250 nL of AAV8-hSyn-DIO-hM3d(Gq)-mCherry (Addgene, Catalog # 44361-AAV8) at a rate of 50 nL/min using a 1000 nL Hamilton syringe bilaterally in the dorsal striatum (coordinates: +1.0 mm and +/- 1.5 mm lateral to Bregma, 2.7 mm deep to the cortical surface). Following 21 days of recovery and allowance for excitatory Gq receptor expression, mice underwent a 14-day tail suspension and administration protocol of clozapine-N-oxide (CNO) (water-soluble, HelloBio, Catalog# HB6149). On Day 0, mice were habituated to handling and the tail suspension procedure. Mice were assessed for clinically-relevant dystonic behavior during tail suspension on Days 1, 7, and 14 as described above. On Day 1, mice were injected i.p. with 1 mg/kg of CNO 30 minutes prior to tail suspension. Mice were then given CNO continuously via their drinking water through Day 14 behavioral analysis. Noting that mouse weights in our cohort at this age ranged between 28-35 g, and given that C57Bl/6J mice drink approximately 8 mL of water / 30 g of body weight per day,(*54*) drinking water solution was made with 0.012 mg of CNO / mL of water to approximate 1 mg/kg/day of CNO delivered to each mouse via drinking water. On Days 7 and 14, mice were again given 1 mg/kg i.p. injection of CNO 30 minutes before tail suspension, to ensure comparability in exposures between each day of the behavioral paradigm (i.e., injections are received by the mice before tail suspension on each day) and to ensure that a full day’s dose of CNO was delivered to these animals on the day of behavior regardless of whether they had all consumed the same amount of water on that day.

### Mouse brain slice preparation and cell-attached recordings

To identify ChIs in brain slices, striatal ChI firing rates were compared between groups of ChAT-Cre positive mice injected with either AAV8-hSyn-DIO-hM3d(Gq)-mCherry (ChIs would express mCherry and the Gq receptor and be excited by CNO) or injected with AAV8-hSyn-DIO-mCherry (Addgene Catalog # 50459-AAV8, ChIs would express only mCherry and not be excited by CNO) as described above. All mice underwent the same CNO exposure paradigm as mice undergoing behavioral assessment. Striatal ChI firing rates were assessed in slice preparations on Day 1 and Day 14 of this CNO exposure paradigm, at least 6 hours after the last exposure to CNO.

For slice preparation, mice were anesthetized with isoflurane and decapitated according to protocols approved by the Washington University Institutional Animal Care and Use Committee. The brain was removed from the skull and attached to a Leica VT1200 specimen holder. Coronal brain slices (300 μm thick) were cut in ice-cold, modified NMDG-HEPES recovery aCSF (in mM: 92 NMDG, 2.5 KCl, 1.25 NaH_2_PO4, 30 NaHCO3, 20 HEPES, 25 glucose, 2 thiourea, 5 Na-ascorbate, 3 Na-pyruvate, 0.5 CaCl_2_, and 10 MgSO_4_; 300 mOsm; pH 7.3–7.4). Slices were transferred into modified NMDG-HEPES recovery aCSF at 32° C. A Na^+^-rich spike-in solution (4 ml, 2 M) was added to gradually increase Na^+^ concentration to improve the success of gigaseal formation. After recovery, slices were kept in modified HEPES holding aCSF (in mM: 92 NaCl, 2.5 KCl, 1.25 NaH_2_PO_4_, 30 NaHCO_3_, 20 HEPES, 25 glucose, 2 thiourea, 5 Na-ascorbate, 3 Na-pyruvate, 2 CaCl_2_, and 2 MgSO_4_; 300 mOsm; pH 7.3-7.4) for at least 1 hour at room temperature before experimental recording.

For recording, slices were transferred to a recording chamber with continuous perfusion (2 ml/min, 32° C) of oxygenated, Na^+^-rich aCSF (in mM: 125 NaCl, 25 glucose, 25 NaHCO_3_, 2.5 KCl, 1.25 NaH_2_PO_4_, equilibrated with 95% oxygen-5% CO_2_ plus 2 CaCl_2_, and 1 MgCl_2_, 310 mOsM). Target interneurons were labeled and identified by fluorescence. IR-DIC microscopy (Nikon FN1 microscope and Photometrics Prime camera) was used to visualize fluorescent cells for pipette attachment. A MultiClamp 700B amplifier, Digidata 1550 16-bit A/D converter, and pClamp 10.4 software (Molecular Devices) were used for recordings. To form a loose-seal, cell-attached patch, Borosilicate glass pipettes (World Precision Instruments, Inc), filled with regular aCSF, had an open-tip resistance of 3-6 MΩ. Action potential activity was recorded in current-clamp mode with sampling rate at 20 kHz and filtered at 6 kHz.

### Mouse brain immunohistochemistry and quantification

Mice were deeply anesthetized with isoflurane and transcardially perfused with 1X PBS followed by 4% paraformaldehyde solution. Extracted brains were placed in 4% PFA solution overnight and were stored in 30% sucrose until further processing. Prior to slicing and staining, brains were treated overnight with 20% glycerol and 2% dimethylsulfoxide to prevent freeze-artifacts. The brains were rapidly frozen, after curing by immersion in 2-Methylbutane chilled with crushed dry ice and mounted on a freezing stage of an AO 860 sliding microtome and sliced into 20μM coronal sections collected in Antigen Preserve solution (50% PBS pH7.0, 50% Ethylene Glycol, 1% Polyvinyl Pyrrolidone); no sections were discarded. For immunofluorescent histochemistry, free floating sections were stained for ChAT (Primary: ChAT Fluoro Stain, Millipore ab144p; Secondary: Anti-Goat AlexaFluor 647, Jackson Labs 705-605-147) and mCherry (Primary: mCherry Double Stain - ChAT/f, EnCor RPCA-mCherry; Secondary: Anti-Rabbit Cy3, Jackson Labs 711-165-147). All incubation solutions from the primary antibody onward use Tris buffered saline (TBS) with Triton X-100 as the vehicle; all rinses are with TBS. The sections were immunostained with the primary antibodies overnight at room temperature.

Vehicle solutions contained Triton X-100 for permeabilization. Following rinses, a fluoro-tagged secondary antibody (anti IgG of host animal in which the primary antibody was produced) was applied. Following rinses, for sections which a biotinylated secondary antibody was applied a fluorescent tagged streptavidin was applied. Following rinses, Hoechst (a nissl counterstain) was applied. Following further rinses, the sections were mounted on gelatin coated glass slides, air dried. The slides were dehydrated in alcohols, cleared in xylene and coverslipped.

### Striatal DREADD expression quantification

To determine the extent of DREADD expression in striatal ChIs, mounted sections were imaged using a Keyence BZ-X810 fluorescence microscope (Keyence Corporation of America, Itasca, IL). For each slice ranging from 145-945 µm anterior to bregma, each panel containing striatum was imaged with a 10X objective using the Multi-Color setting to capture an overlay of ChAT and mCherry co-expression. Built-in autofocus software was used on each panel to ensure optimal image quality. Panels were combined using Image Stitch software in the BZ-X800 Analyzer to generate a full, uncompressed image of each striatal section. Although exposure settings were consistent across panels within each slice, exposure varied across slices and across animals. Imaging was primarily used for particle counting, a metric that checks for presence or absence rather than fluorophore intensity. We quantified DREADD expression in each striatal section by using ImageJ to measure ChAT and mCherry co-expression relative to the total population of striatal ChIs. To count the total number of ChIs in each striatal section, we converted all images to Grayscale (8-bit) with a minimum default threshold of 10. We processed the resulting selection using watershed segmentation and then used the built-in Analyze Particles function, setting the minimum particle size to 9 pixels, to count the total number of selected neurons within the striatum region within the slice. To determine the number DREADD-expressing ChIs in each striatal section, the neurons expressing both ChAT and mCherry were manually counted in each slice and cross-referenced to neurons identified with automated counting to ensure any neurons that were manually identified were also included in the total calculated ChI population. For any given slice, the percentage striatal ChIs expressing the DREADD was calculated by dividing the number of mCherry-expressing ChIs by the total number of ChIs.

### Statistical Analyses

Statistical analysis was done in GraphPad Prism (version 8, GraphPad Software), SPSS (version 28.0, IBM), or within an Anaconda virtual environment with the following packages feature selection analyses; python 3.9.16, scikit-learn 1.2.2, scipy 1.10.0 and mrmr-selection 0.2.6. Leg adduction variability and amplitude metrics in human subjects were compared between videos with and without expert-identified dystonia using t-tests when appropriate (with normality of data sets assessed using the Shapiro-Wilk test). For data with unequal standard deviations between groups, a t-test with Welch’s correction was applied. For data that were not normally distributed, Mann-Whitney tests were used. Pearson correlations and multiple linear regression were used to determine whether these leg adduction metrics were significantly correlated to expert assessments of dystonia severity. For determining which variability and amplitude metrics best discriminate between the presence or absence of dystonia or predicted dystonia severity in human subjects, 7 filter, wrapper, and embedded feature selection methods were used (Spearmans rho, Kendalls tau, univariate feature selection, maximum relevance - minimum redundancy, extra trees classifier, random forest, and sequential feature selection (forward)). The methods were chosen based on their widespread use for feature selection and ability to handle non-normal distributions, as well as their suitability for ratio variable types and distance metrics.(*55*) These methods use different decision values, such as p-value and classification accuracy, for selecting the metrics that best discriminate between the presence or absence of dystonia or predict dystonia severity. The top 4 metrics out of 8 distance and variability metrics were selected using each of the 7 feature selection methods. Two-way repeated measures ANOVAs were used to determine whether there were significant differences in leg adduction variability and amplitude metrics in response to acute vs. chronic CNO administration. T-tests and paired t-tests were used to assess differences in in striatal ChI firing rate between mice either expression or not expressing DREADDs (t-test) or in cells before and after CNO (paired t-test). The significance level for all tests was set *a priori* to p<0.05.

## List of Supplementary Materials

(also available here: https://wustl.box.com/s/07agml8jmw681et1bnsk2j4atghy5j9l)

Materials and Methods S1. MATLAB code used to calculate leg adduction variability and amplitude in mice. (Chronic_Striatal_ChI_Dystonia_Mice.m)

Materials and Methods S2. MATLAB code used to calculate leg adduction variability and amplitude in people. (Dystonia_CP_Seated_Human.m)

Figure S1. Correlation between leg dystonia severity and leg adduction amplitude and variability in people with CP.

Figure S2. Leg adduction amplitude and variability in mice in response to 14 days of striatal ChI excitation.

Table S1. Demographic characteristics of people with CP assessed for dystonia.

Table S2. The top leg adduction metrics contributing to models discriminating between the presence versus absence of dystonia or to models predicting dystonia severity.

Data file S1. Leg adduction amplitude and variability values in people with CP (Sheet 1), leg adduction amplitude and variability values in mice from 1 to 14 days of striatal ChI excitation (Sheet 2), striatal ChI DREADD expression (Sheet 3), striatal ChI firing rates (Sheet 4) (Data_Chronic_Striatal_ChI_Dystonia.xls)

Movie S1. De-identified video of a person with CP without leg dystonia (median GDRS 0) labeled with DeepLabCut (GDRS 0 - blurred labeled cropped.mp4)

Movie S2. De-identified video of a person with CP with mild leg dystonia (median GDRS 3) labeled with DeepLabCut (GDRS 3 - blurred labeled cropped.mp4)

Movie S3. De-identified video of a person with CP with moderate leg dystonia (median GDRS 13.5) labeled with DeepLabCut (GDRS 13.5 - blurred labeled cropped.mp4)

Movie S4. Video of a control mouse following 14 days of CNO administration (no striatal ChI excitation) labeled with DeepLabCut (Control_37M2_labeled.mp4)

Movie S5. Video of an experimental group mouse following 14 days of CNO administration (chronic striatal ChI excitation) labeled with DeepLabCut (Chronic_ChI_Excitation_53F1_labeled.mp4)

---

**Figure S1.**
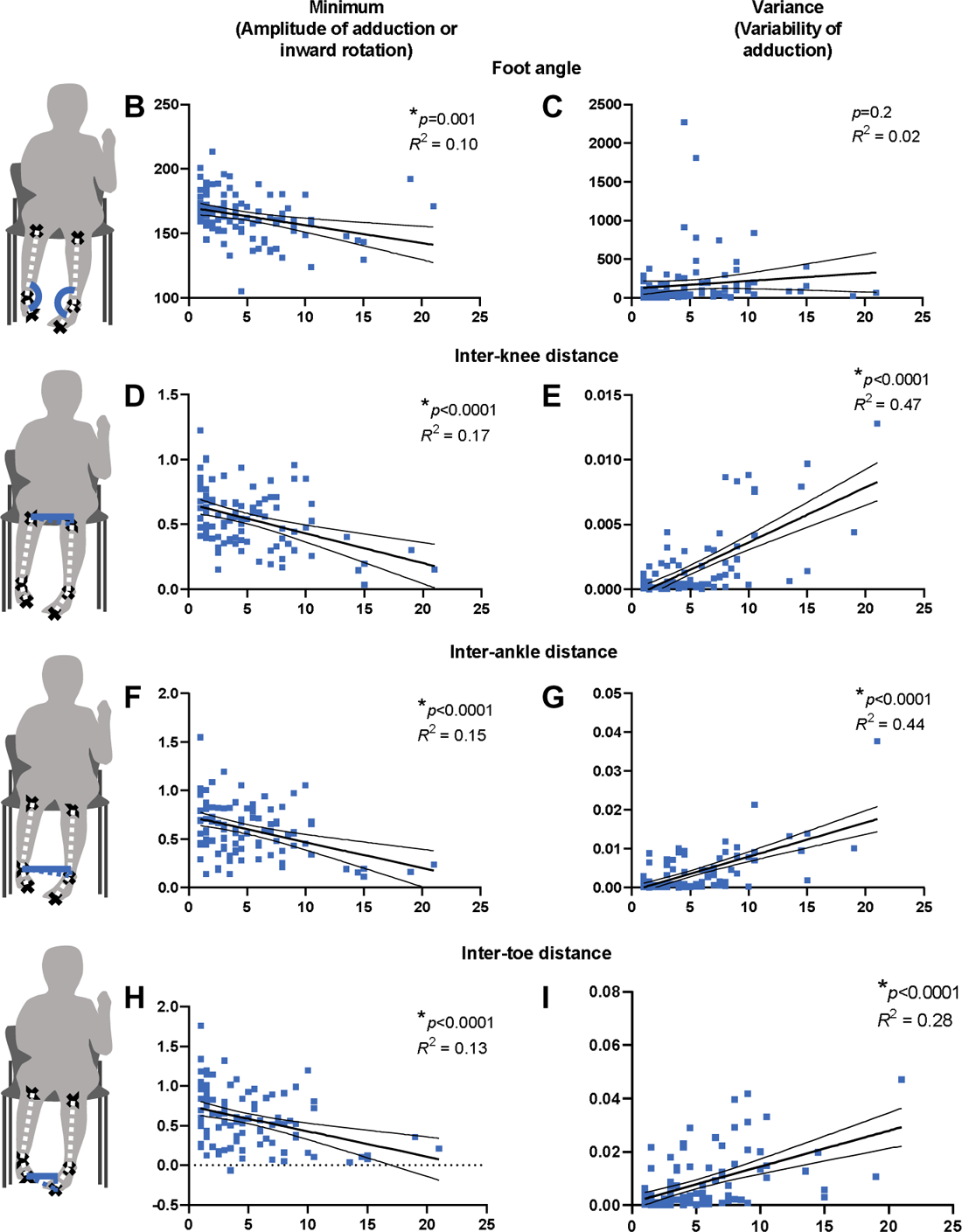
Correlation between leg dystonia severity and leg adduction amplitude and variability in people with CP. Metrics of leg adduction amplitude (left column) and variability (right column) are correlated with average Global Dystonia Rating Scale (GDRS) scores indicating expert-assessed leg dystonia severity in people with CP (blue, n=103). Metrics are the minima (left column) and variances (right column) of the foot angle in the coronal plane (**A&B**: angle between the knee, ankle, and toe labels), inter-knee distance (**C&D**), inter-ankle distance (**E&F**), and inter-toe distance (**G&H**) with distance metrics assessed exclusively along an X-axis parallel to the floor. Pearson correlation, *p<0.05.

**Figure S2.**
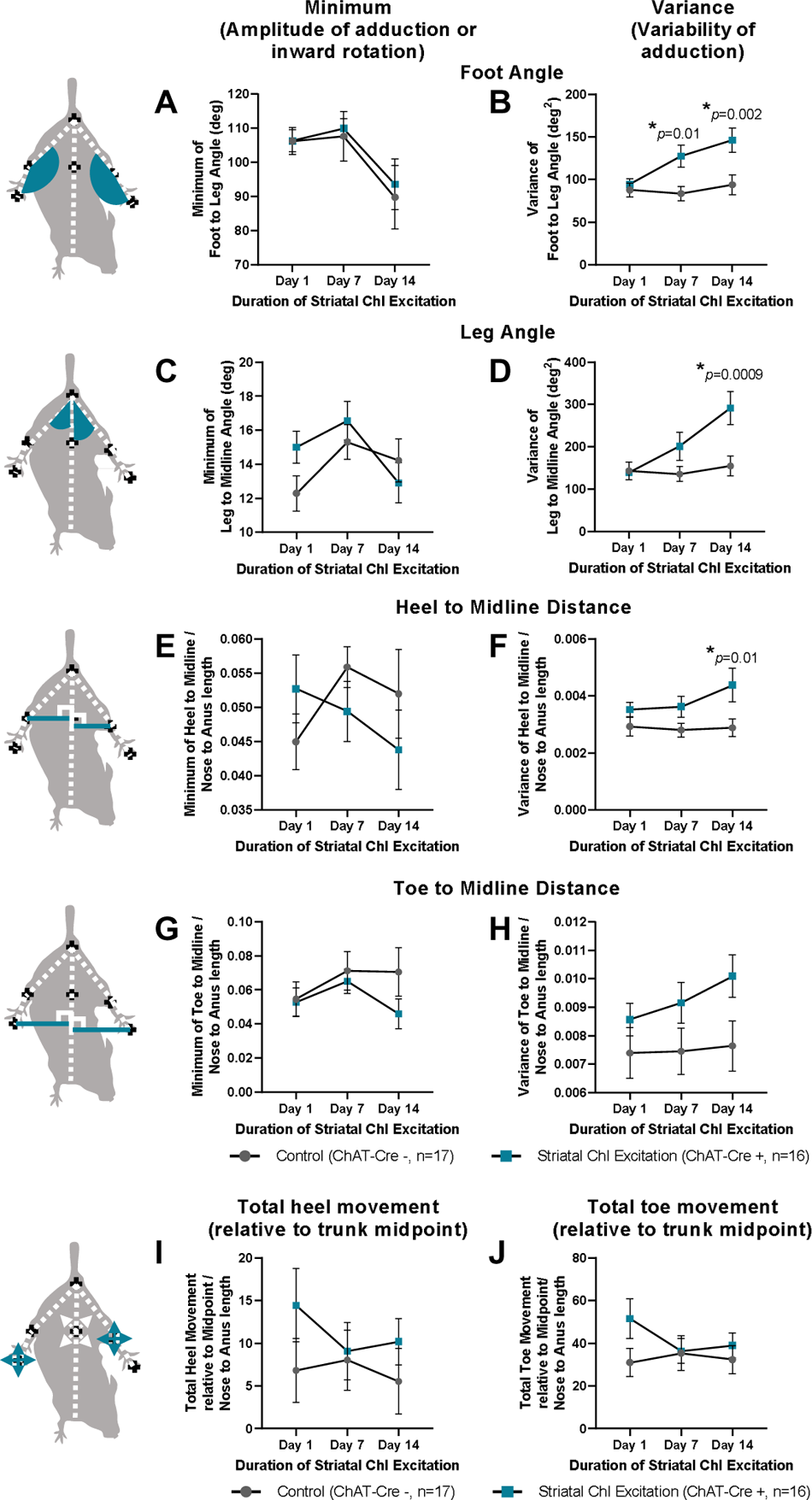
Leg adduction amplitude and variability in mice in response to 14 days of striatal ChI excitation. **A-H)** Leg adduction metrics are the amplitudes or minima (left column) and variability or variances (right column) of the foot angle between the hindpaws and legs (**A&B**), leg angle between the leg and the mouse midline (**C&D**), distance between the heels and midline (**E&F**), and distance between the toes and midline (**G&H**). **I&J)** Total movement of the heels (**I**) and toes (**J**) relative to the mouse midpoint. Comparisons between mice that did and did not undergo ChI excitation are by repeated measures two-way ANOVA with Bonferroni correction for multiple comparisons, *p<0.05.

**Table S1.**
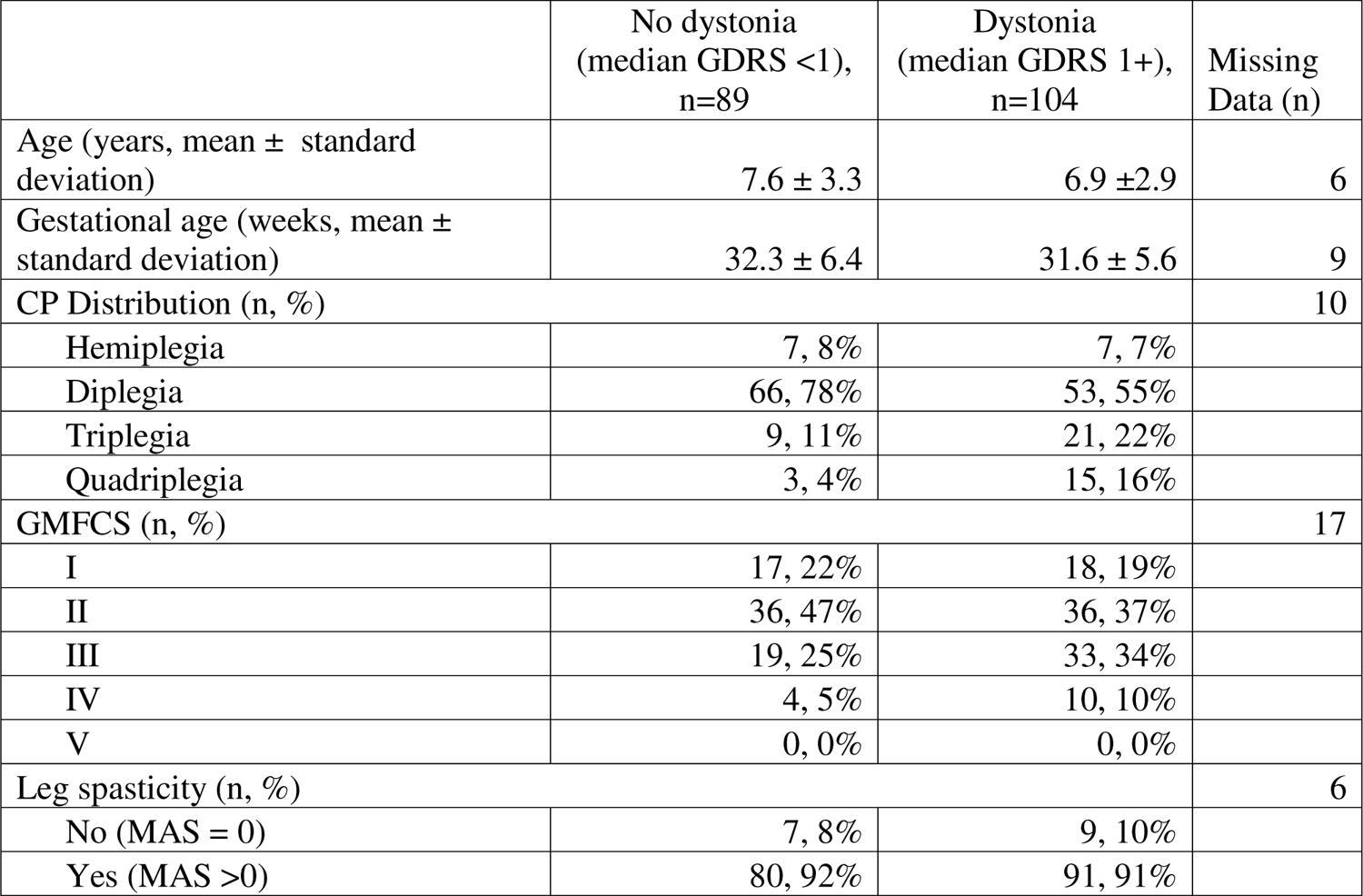
Demographic characteristics of people with CP assessed for dystonia. GDRS – Global dystonia rating scale, with median score determined between 8 pediatric movement disorders specialists rating leg dystonia. GMFCS – Gross Motor Functional Classification System (I-II – independently ambulatory without assistive devices, III – independently ambulatory with assistive devices, IV-V – uses wheelchair for mobility). MAS – Modified Ashworth Scale (a measure of spasticity, with 0 indicating no spasticity and scores of 1-3 indicating increasing severities of spasticity).

**Table S2.** The top leg adduction metrics contributing to models discriminating between the presence versus absence of dystonia or to models predicting dystonia severity. Seven models (left) were used to identify the top 4 metrics out of 8 total (4 variability + 4 amplitude metrics) that could discriminate between the presence or absence of leg dystonia (middle) or predict leg dystonia severity (right). The number of variability or amplitude metrics that were selected as the top 4 metrics in each model is indicated.

| Feature selection method | Models discriminating between presence vs. absence of dystonia |  | Models predicting dystonia severity |  |
| --- | --- | --- | --- | --- |
|  | Amplitude | Variability | Amplitude | Variability |
| Spearman's rho | 1 | 3 | 1 | 3 |
| Kendall's tau | 1 | 3 | 1 | 3 |
| Anova F-test /univariate feature selection | 1 | 3 | 1 | 3 |
| Maximum Relevance - Minimum Redundancy | 1 | 3 | 0 | 4 |
| ExtraTreesClassifier | 1 | 3 | - | - |
| Random forest | 1 | 3 | - | - |
| Sequential Feature Selection (forward) | 0 | 4 | - | - |

## Funding

BRA – NINDS K08NS117850, Pediatric Epilepsy Research Foundation

JGM – NINDS R01NS117899, Rita Allen Foundation

MCK – NINDS R01NS106298 SM – NIMH R01MH123748

## Author contributions

Conceptualization: BRA, JGM Methodology: BRA, JGM, SM

Investigation: BRA, KG, XL, KC, AR, RB, NS, JB, RGM, MCK, DM, JOM, LT, JW, SW, TF

Visualization: BRA, KG, XW, RB

Funding acquisition: BRA, JGM, SM

Project administration: BRA, KC

Supervision: BRA, JGM

Writing – original draft: BRA

Writing – review & editing: BRA, JGM, SM, JP, XL, KG

## Data and materials availability

All data are available in the main text or the supplementary materials.

